# A synthetic, three-dimensional bone marrow hydrogel

**DOI:** 10.1101/275842

**Authors:** Lauren E. Jansen, Thomas P. McCarthy, Michael J. Lee, Shelly R. Peyton

## Abstract

Three-dimensional (3D) synthetic hydrogels have recently emerged as desirable *in vitro* cell culture platforms capable of representing the extracellular geometry, elasticity, and water content of tissue in a tunable fashion. However, they are critically limited in their biological functionality. Hydrogels are typically decorated with a scant 1-3 peptide moieties to direct cell behavior, which vastly underrepresents the proteins found in the extracellular matrix (ECM) of real tissues. Further, peptides chosen are ubiquitous in ECM, and are not derived from specific proteins. We developed an approach to incorporate the protein complexity of specific tissues into the design of biomaterials, and created a hydrogel with the elasticity of marrow, and 20 marrow-specific cell-instructive peptides. Compared to generic PEG hydrogels, our marrow-inspired hydrogel improves stem cell differentiation and proliferation. We propose this tissue-centric approach as the next generation of 3D hydrogel design for applications in tissue engineering.

---

The vast majority of materials available to study how environmental cues direct cell fate are two-dimensional (2D), ranging from protein-coated surfaces to hydrogels^1-3^. However, 2D materials restrict cell adhesions to an x-y plane and force an apical-basal polarity^4,5^. To overcome this, researchers can better recapitulate the *in vivo* geometry of tissues using hydrogels to culture cells in three-dimensions (3D)^6^. Synthetic hydrogels made with polyethylene glycol (PEG) can be functionalized with peptide motifs that either elicit integrin-binding or allow for cell-mediated matrix degradation. Additionally, PEG hydrogels are extremely reproducible and are independently tunable in both stiffness and ligand density. Nevertheless, in some instances PEG-based hydrogels are generally considered inferior to more complex protein-based hydrogels, like Matrigel, which may more accurately mimic native tissue. A main limitation in the use of naturally derived protein hydrogels is that the constituents are undefined and have extreme batch-to-batch variability. Thus, there is a need for synthetically engineered hydrogels designed to mimic structural and physiochemical features of specific *in vivo* environments.

Bone marrow is the soft interior tissue between hard compact bone where many of our immune and stromal stem cells reside. Like every human tissues and organ, bone marrow has unique biophysical features that are critical for cell and organ function. For example, protein composition and tissue stiffness are important for cellular processes like migration and proliferation^7,8,9^, as well as regulating stem cell fate and organoid development^10-12^. Thus, it is not surprising that the surrounding extracellular matrix (ECM) plays a key role in the proper function of bone marrow because both hematopoietic and stromal progenitor cells originate from this tissue^13^. For example, both bone marrow stiffness and fibronectin are important for the maintenance of hematopoietic stem cell progenitors^14^. Additionally, marrow-derived stromal stem cells will differentiate into either bone or fat cells in response to mechanical cues^15^ and the presence or absence of vitronectin in 3D scaffolds can facilitate reversible differentiation into or from osteoblasts^16^.

Despite clear evidence of the marrow ECM regulating the stem cell niche, *in vitro* stem cell culture platforms contain a mere fraction of the biochemical cues typical of the native tissue. Simple RGD-decorated polymers do not fully capture these cues, making it imperative to move toward environments that include the diversity of integrin-binding and protease-sensitive proteins in native tissues and organs. We propose a 3D ECM-focused material based on PEG and peptides. Unique to our approach is the development of a combination of bioinformatics and biomechanics to create a set of tissue-specific design criteria. This new class of tissue-centric PEG hydrogels captures the protein complexity of the native tissue in a material that is extremely tunable and can be fabricated with little technical expertise.

## A biomechanics and bioinformatics approach to create a synthetic human bone marrow

A top-down engineering approach was used to identify the physical and chemical properties of bone marrow that could be represented by a synthetic PEG hydrogel (Figure 1a-b). Bone marrow elasticity was measured via shear rheology, indentation, and cavitation rheology^7^. We then approximated this elasticity synthetically by adjusting the polymer-polymer distance of the hydrogel, a material that is inherently hydrophilic and mimics marrow’s high water content. We have previously demonstrated that bone marrow is a benign elastic material^7^. Because the viscoelastic properties contribute to stem cell fate^17^, we compared the compressive properties of both porcine marrow and of a PEG hydrogel. Both materials closely followed a Hertzian model (Suppl. Figure 1 a-b), suggesting that under these conditions, PEG is an appropriate physical model for the elasticity of marrow.

**Figure 1.**
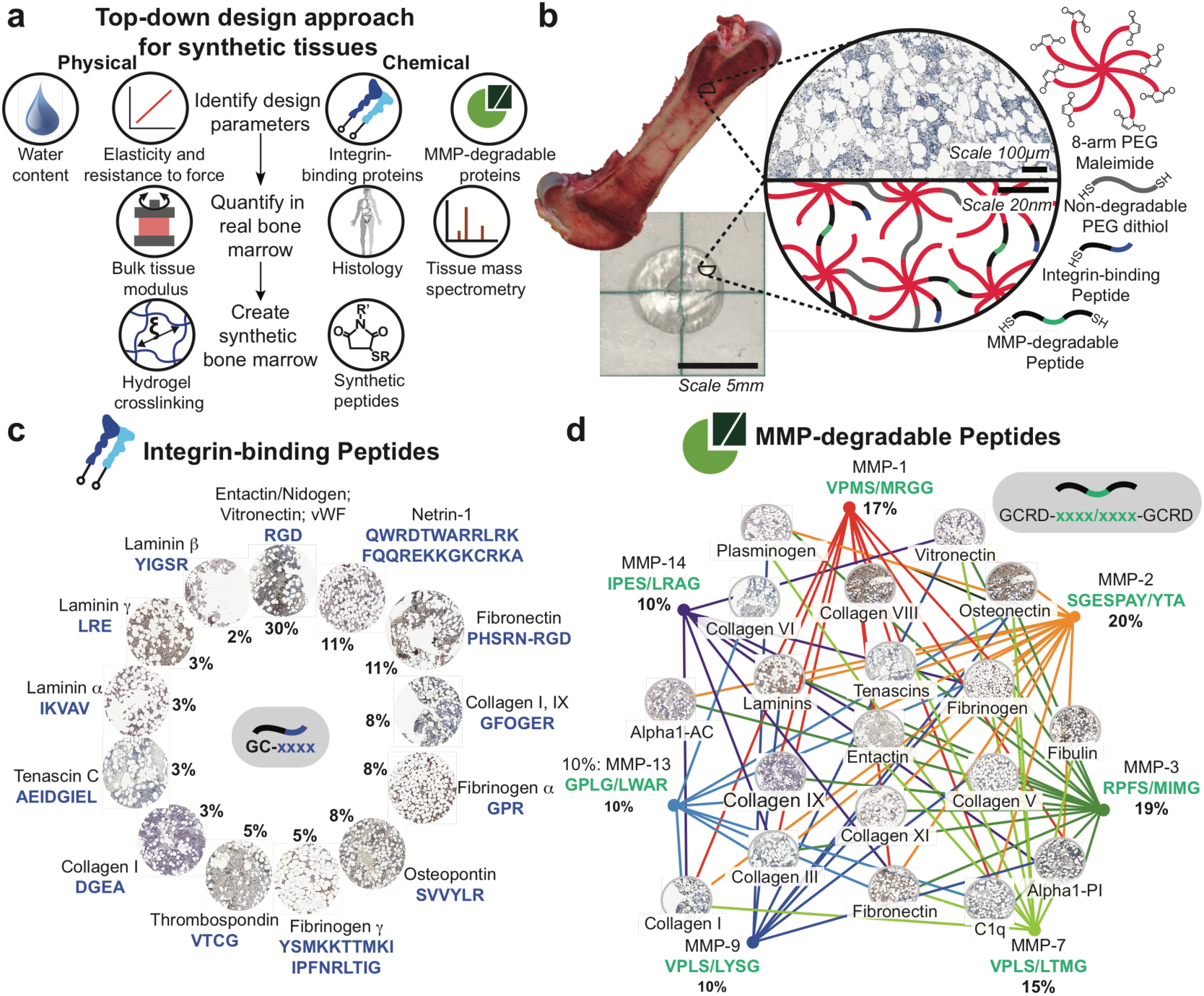
Adapting a PEG hydrogel to mimic the physical and chemical properties of bone marrow tissue. a) Tissues have specific defining physical and chemical properties such as water content, elasticity, integrin-binding, and MMP-degradable proteins. These properties can be quantified in real bone marrow tissue using rheology, mass spectrometry, and tissue histology (body image from Protein atlas). These features are represented synthetically by tuning the hydrogel crosslinking and incorporating biofunctional peptides. c) Here, bone marrow tissue (top left, image from Jansen et al., 2015) is mimicked in a hydrogel (bottom left) composed of an 8-arm PEG macromer functionalized with c) 13 mono-functional integrin-binding peptides, and crosslinked with d) 7 di-functional MMP-degradable peptides and a linear PEG-dithiol. The known functional sequence for each peptide is depicted in blue with the percentage it is present in the hydrogel (% relative to other peptides). All histology images are representative of each protein in human bone marrow tissue (images from the Protein Atlas). The lines in d) connect each MMP to their known protein substrates and the slash (/) indicates the cleavage location for each enzyme on the matched peptide.

The ECM proteins of marrow were identified using histology and mass spectrometry (Figure 1a). ECM proteins were represented by specific peptide sequences that are either responsible for high affinity binding to transmembrane integrin proteins ^18^ or are highly susceptible to cleavage by matrix metalloproteinases (MMPs) (Suppl. Table 1 and 2). Integrins are the largest class of cell adhesion receptors that mediate attachment to the ECM and activate intracellular signaling^19^. Collectively the MMP family can degrade all components of the ECM and is important for tissue homeostasis^20^. Integrin-binding proteins and MMP protein substrates were identified in human bone marrow using the histology data available in the Protein Atlas (Suppl. Table 3)^8^. Peptide motifs that elicit integrin binding were identified for each protein and displayed as mono-functional moieties in the hydrogel (Figure 1b-c, Suppl. Table 5-6)^21-29^. Conversely, di-functional peptides that selectively degrade in the presence of cell secreted MMPs were associated with the protein substrates found in marrow (Figure 1b,d, Suppl. Table 6). The histological scores for each protein were used to determine the quantitative peptide amounts to be included in the final bone marrow hydrogel (Figure 1c-d). Representative images from the tissue histology in the Protein Atlas are displayed in Figure 1c and d. ECM proteins in human bone marrow^30^ were analyzed via liquid chromatography-mass spectrometry (LC-MS, Suppl. Figure 1c) and compared to control tissues (lung and brain, Suppl. Table 4) to confirm that our histology-driven approach identified the unique ECM signature of marrow. The LC-MS data from human bone marrow was most similar to the bone marrow peptide cocktail (Suppl. Figure 1d), and protein substrates from bone marrow ECM could be cleaved by active MMP enzymes (Suppl. Figure 1e). Together, these data indicated our technique could accurately filter for the unique integrin-binding and MMP-degradable protein signature of marrow.

## Functional validation of peptide domains

Human mesenchymal stem cells (MSCs) were used to test whether stromal cells, which are highly abundant in the marrow, could recognize and respond appropriately to the peptides in our bone marrow cocktail. We developed a competitive cell adhesion assay to measure binding to integrin peptides, which took advantage of the decrease in cell area observed during cell adhesion (Suppl. Video 1 and 2). We covalently attached the full integrin-binding peptide cocktail (Figure 1c) to a glass coverslip upon which MSCs were seeded in the presence or absence of individual peptides from that same cocktail solubilized in the cell medium (Figure 2a). We validated that cell adhesion to the surface was driven by the attached peptides, not serum (Suppl. Figure 2a), and that protein did not passively bind to peptide-coated coverslips in the experimental time frame (Suppl. Figure 2b). We observed a decrease in the area of cells pre-treated with peptides, so adhesion was quantified using cell area measurements at two hours (Figure 2a).

**Figure 2.**
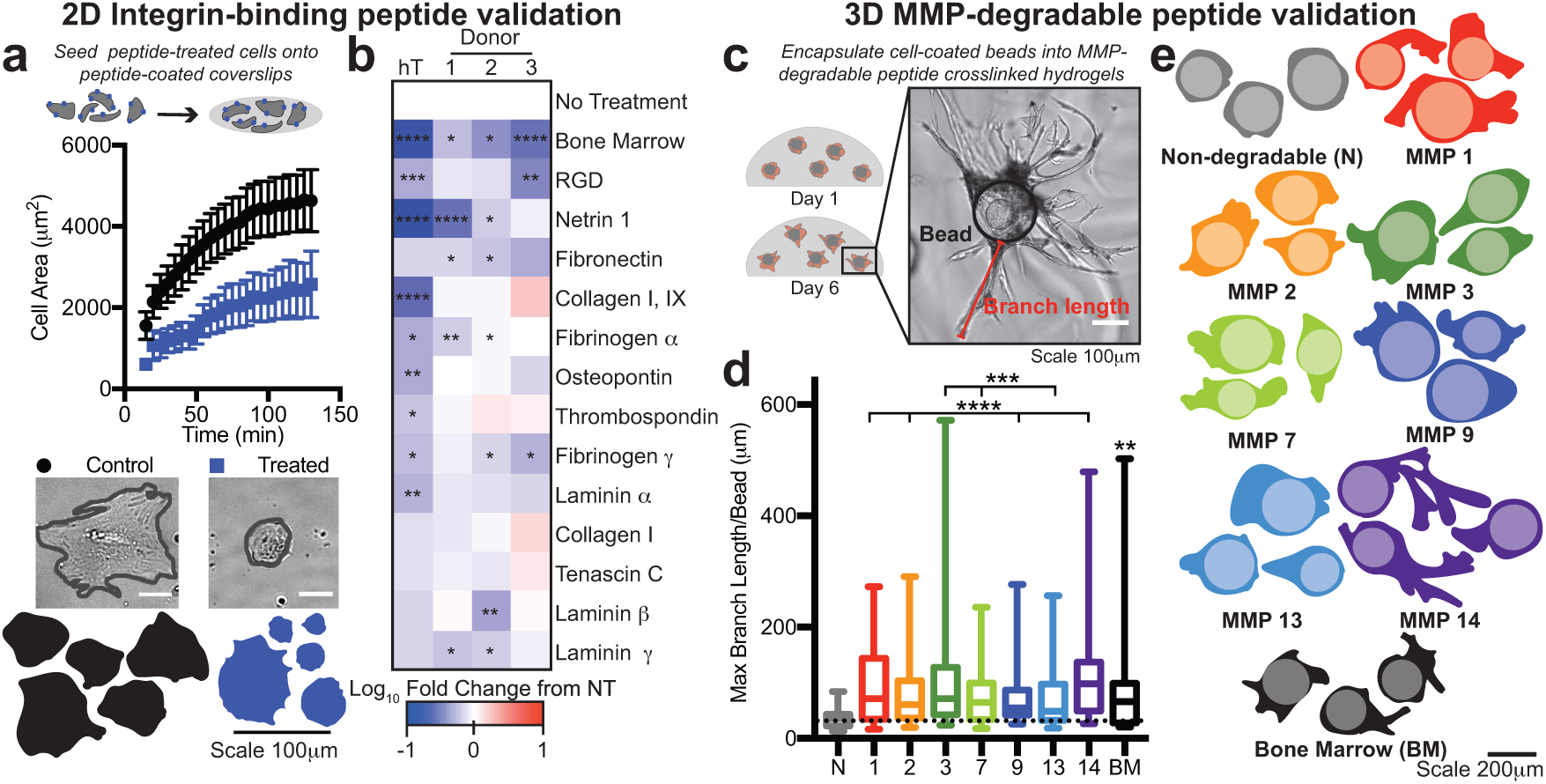
Peptide moieties can be bound and degraded by MSCS. a) Cells were treated with peptides and seeded onto coverslips coated with the bone marrow peptide cocktail. MSC area was measured over time for cells not treated (control, black) or pre-treated for 30 minutes prior (blue) with soluble integrin binding peptides and allowed to adhere to a surface coupled with the bone marrow peptide cocktail. Representative cell images (scale 50 μm) and outlines of MSCs 2 hours after seeding (bottom). Error bars represent SEM. b) Heat map depicting the log10 fold change in cell area at 2 hours compared to no treatment (NT) for each integrin-binding peptide moiety in the mimic across one cell line, hTERT MSCs (hT), and three donor MSCs (1-3) (BM=bone marrow peptide cocktail) (N≥2, n≥20 per cell). c) Representative image of MSCs seeded on cytodex beads (black outline) and encapsulated into a hydrogel with one MMP degradable crosslinker (Cell area=red, branch length=green). d) A box and whisker plot for the maximum branch length per bead in each hydrogel condition. e) Representative cell and bead traces in each hydrogel condition, where the lighter colored circle is the bead and the darker color is the cell trace (N=2, n≥15 per cell). Significance is

All tested cells had decreased adhesivity when dosed with the bone marrow integrin-binding peptide cocktail (Figure 2b, Suppl. Figure 2d). Across three MSC sources from human donors and one immortalized cell line, most individual peptides decreased MSC spreading at the concentration at which they were present in the cocktail (Figure 2b). The Collagen I and Tenascin C peptides did not significantly alter MSC adhesion in any case, but promoted an anti-adhesive phenotype in a human breast cancer cell line (Suppl. Figure 2d). Interestingly, the immortalized MSC cell line was more responsive to individual peptide treatments compared to the donor cells (i.e. more peptides decreased adhesion). Since ECM proteins can facilitate both adhesion to and dissociation from the matrix, and we see those phenotypes in this data, we conclude that each peptide in the mimic can direct cell adhesion to and from our hydrogel matrix.

Crosslinker degradation was validated using a cell invasion assay. Cytodex beads were coated with MSCs and encapsulated for 6 days into the hydrogels crosslinked with a single MMP degradable peptide or the full set of degradable crosslinks at the same molar ratio of thiol to maleimide (Figure 2c). When degradable peptides were present, MSCs were able to branch further into the surrounding network (Figure 2d-e). MSCs branched the furthest in the bone marrow-cocktail, MMP-3, and -14 crosslinked hydrogels, suggesting that certain individual peptides can be extremely susceptible to degradation and peptide combinations can improve material degradation by bone marrow cells. Since all the MMP peptides have been optimized to selectively degrade in response to their respective enzyme^22^, we suggest that MMP expression would likely explain the differential degradation observed. In sum, we observed degradation for all MMP peptide crosslinkers and found that the full bone marrow-inspired degradable peptide cocktail was optimal for MSC branching.

## Optimal conditions for coupling marrow-specific peptides

This is the first time 20 unique peptide motifs have been incorporated into a PEG hydrogel. Prior to this work, the most complex bioactive PEG hydrogel contained 3 unique peptides^31,32^. Thus, it was important to identify the ideal conditions to incorporate these peptides into the network. Our peptides are coupled using a Michael-type addition reaction, with a maleimide as the Michael-type acceptor. The maleimide-thiol reaction is biocompatible and has been shown to provide the most efficient incorporation of ligands and the largest range of bulk properties compared to similar PEG hydrogels^33^. A thiol quantification assay was used to identify uncoupled peptides in solution because the Michael-type donor for this reaction is a thiol (Suppl. Figure 3a). A number of parameters regulated the efficiency of peptide incorporation, including polymer wt% and the molar percentage of reactive pairs (Suppl. Figure 3b-e). However, these properties also change the effective Young’s modulus of the hydrogel. Overall, we found that an 8-arm PEG enabled increased crosslinking without reducing reaction efficiency (Suppl. Figure 3d).

**Figure 3.**
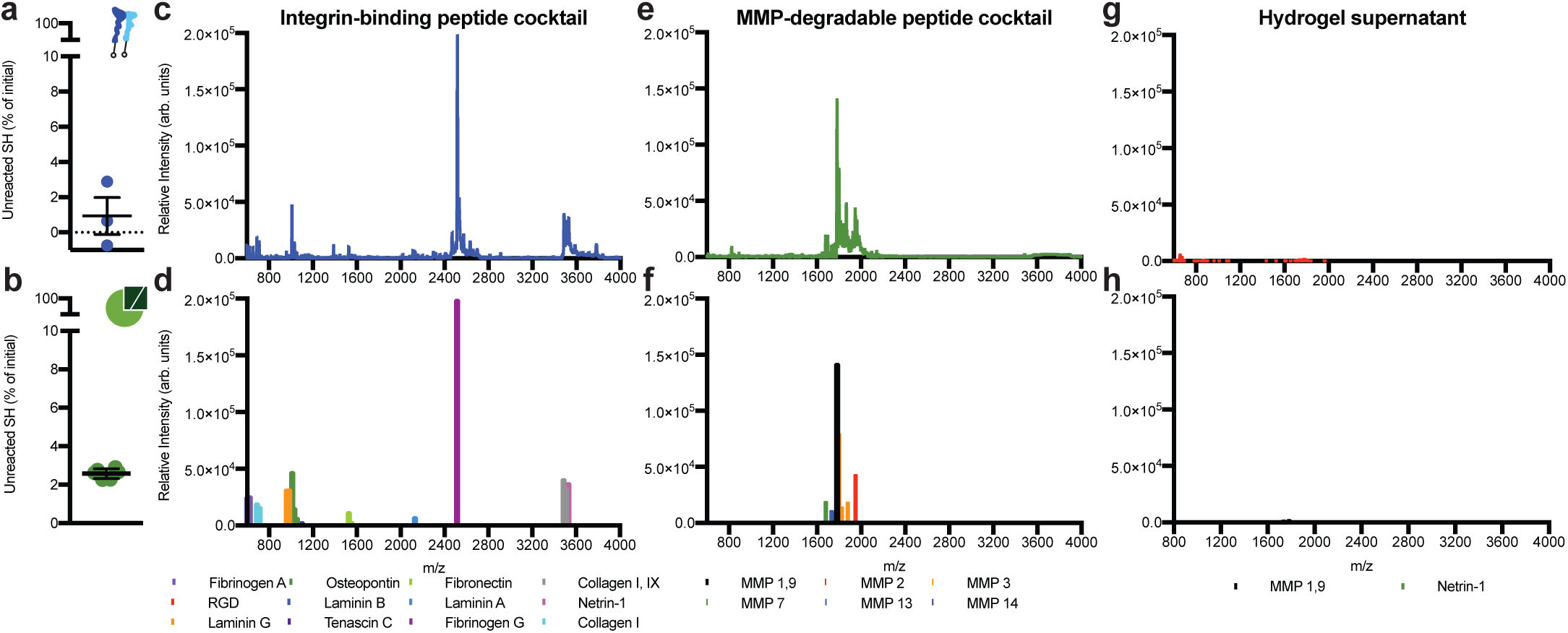
Bone marrow peptides couple to the hydrogel at the expected concentration. a) The percentage of unreacted thiols when mono-functional peptides are added to a solution of PEG dissolved in PBS at pH 7.4. b) The percentage of unreacted thiols 10 minutes post-crosslinking an 8-arm PEG hydrogel at a 1:1 molar ratio of thiol to maleimide. Error bars represent the SEM (N≥1, n≥3). MALDI-TOF spectrum (top) and identified peptide peaks (bottom) for the c) and d) bone marrow mono-functional peptide cocktail, e) and f) the bone marrow di-functional peptide crosslinkers, and g) and h) the supernatant of a bone marrow hydrogel swelled for 4 hours in PBS.

We achieved >98% coupling of mono-functional integrin-binding peptides and >97% of di-functional MMP degradable peptides using an 8-arm PEG at 20 wt% (Figure 3a-b). Optimal reaction conditions for integrin-binding peptides occurred in PBS at pH 7.4, but we did note that the peptide cocktail was less soluble in this buffer than in DMSO (Suppl. Figure 3f-g). Separately, we reduced a hydrogel using sodium borohydride to ensure that disulfide bonds between the thiols were not preventing the Michael-addition reaction. This reduction did not significantly increase the number of free thiols found, indicating that >95% of the material bonds are from the Michael-type addition reaction (Suppl. Figure 3h).

Matrix assisted laser deposition ionization time of flight was used to identify the peptides that did not couple to the matrix. First, we made a solution of all the peptides, without PEG present, and identified all except DGEA and AEIDGIEL (Figure 3e-f, Suppl. Figure 4a-b). These are both highly negatively charged peptides, which do not ionize easily. To confirm this, AEIDGIEL could not be identified even at a high concentration if non-charged peptides were present (Suppl. Figure 4c-f). We then formed hydrogels with all the peptides, and attempted to find unreacted peptides in solution. Only two peptides were identified in the supernatant but at a significantly reduced intensity (Figure 3g-h). Taken together, this data shows that our peptides are crosslinked into the hydrogel close to the expected concentration.

**Figure 4.**
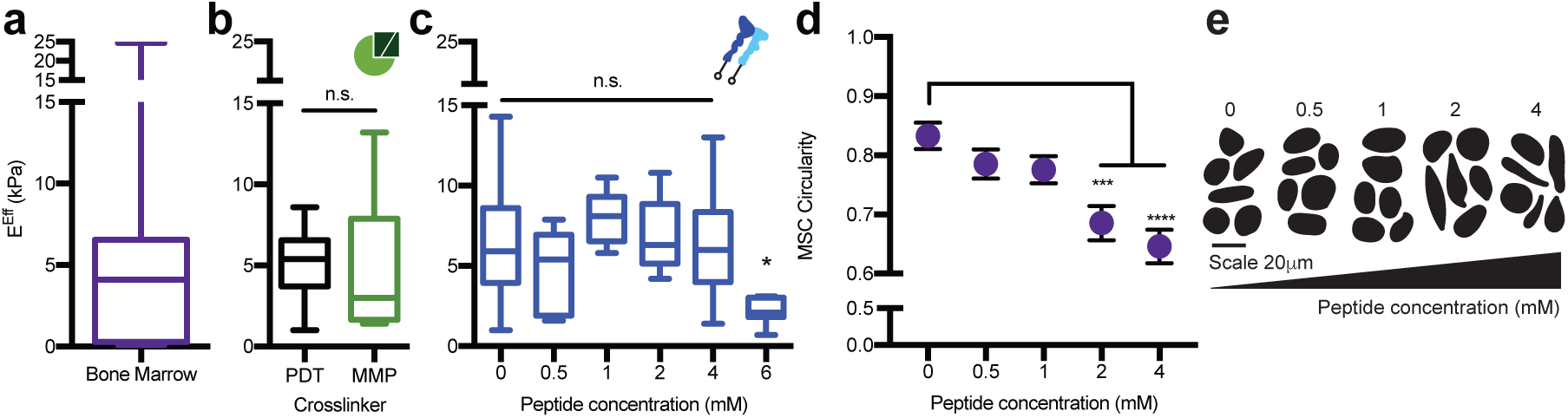
The PEG hydrogel accurately models the bulk compressive properties of bone marrow tissue. a) Rheology data from Jansen et al., 2015 for the effective Young’s modulus (E^Eff^) of porcine bone marrow at 35**°**C. b) The E^Eff^ for 20 wt%, 8-arm, 20K PEG hydrogels crosslinked at a 1:1 thiol to maleimide molar ratio with 1.5K PEG-dithiol (PDT, black) or with the bone marrow cocktail of 1.5K PDT and MMP crosslinkers (MMP, green). c) The E^Eff^ for 20 wt%, 8-arm, 20K PEG hydrogels crosslinked at a 1:1 thiol to maleimide molar ratio with PDT and coupled with different concentrations of the bone marrow peptide cocktail for 10 minutes before gelation. d) MSCs circularity with respect to peptide concentration and e) representative cell traces for cells encapsulated in a 20 wt%, 8-arm, 20K PEG-crosslinked with the bone marrow cocktail. The significance is determined using a two-tailed t-test where p=0.05, and error bars represent the SEM. (N≥2, n≥3 for mechanical testing; N≥2, n≥10 for cell circularity).

## PEG hydrogels mimic the bulk mechanics of bone marrow

The mechanical properties of engineered materials can influence the migration and differentiation of marrow-derived stromal and hematopoietic stem cells^14,15,17,34-37^. We have previously shown that porcine bone marrow has an average modulus of 4.4±1.0 kPa at physiological temperature (Figure 4a)^7^, and recent work has shown that hematopoietic progenitor populations can be maintained in the presence of fibronectin at this modulus^14^. These studies highlight an important role for the mechanical properties of bone marrow tissue to direct stem cell fate and function. A PEG hydrogel crosslinked with a linear dithiol can be adapted to span the range of stiffness observed in bone marrow (Suppl. Figure 3c). While a number of properties contribute to stiffness and can be used to tune modulus, a 20 wt%, 8-arm, 20K PEG hydrogel best matched the modulus of porcine bone marrow tissue (Figure 4b).

A benefit of PEG hydrogels is the mechanical and chemical properties can be independently tuned. To ensure our chemical modifications did not change the stiffness of the material, we individually incorporated them into the hydrogel and tested each of their effects on stiffness. Incorporation of the MMP-sensitive crosslinkers in lieu of PEG-dithiol did not alter the hydrogel modulus (Figure 4b), and mono-functional integrin-binding peptides could be incorporated up to a 4 mM total concentration without compromising the bulk modulus (Figure 4c). Through separate cell tracing experiments, we found that a 2 mM concentration of integrin-binding peptides was needed to achieve significant MSC spreading at 24 hours. We therefore used 2 mM of total integrin-binding peptides in a 20 wt%, 8-arm, 20K PEG crosslinked with our MMP-degradable cocktail as the final bone marrow formulation (Figure 4d-e).

## The synthetic bone marrow hydrogel provides a niche for MSC growth and differentiation

Our results demonstrate an approach to identify the matrix stiffness, integrin-binding peptides, and MMP-degradable sites in real bone marrow. This information was used to create a PEG hydrogel with tissue-inspired properties. As a comparison to the current standard for *in vitro* synthetic 3D culture systems, we compared this bone marrow hydrogel to the commonly used RGD-functionalized PEG hydrogel and tissue culture polystyrene (TCPS). We explored both cell growth and differentiation because these are two phenotypes exhibited by MSCs in real bone. After one week in culture, the same percentage of MSCs expressed Ki67 and p21 on TCPS as in the bone marrow gel, where cells in the RGD-functionalized PEG hydrogel were less proliferative and had increased cellular senescence (Figure 5a-c).

**Figure 5.**
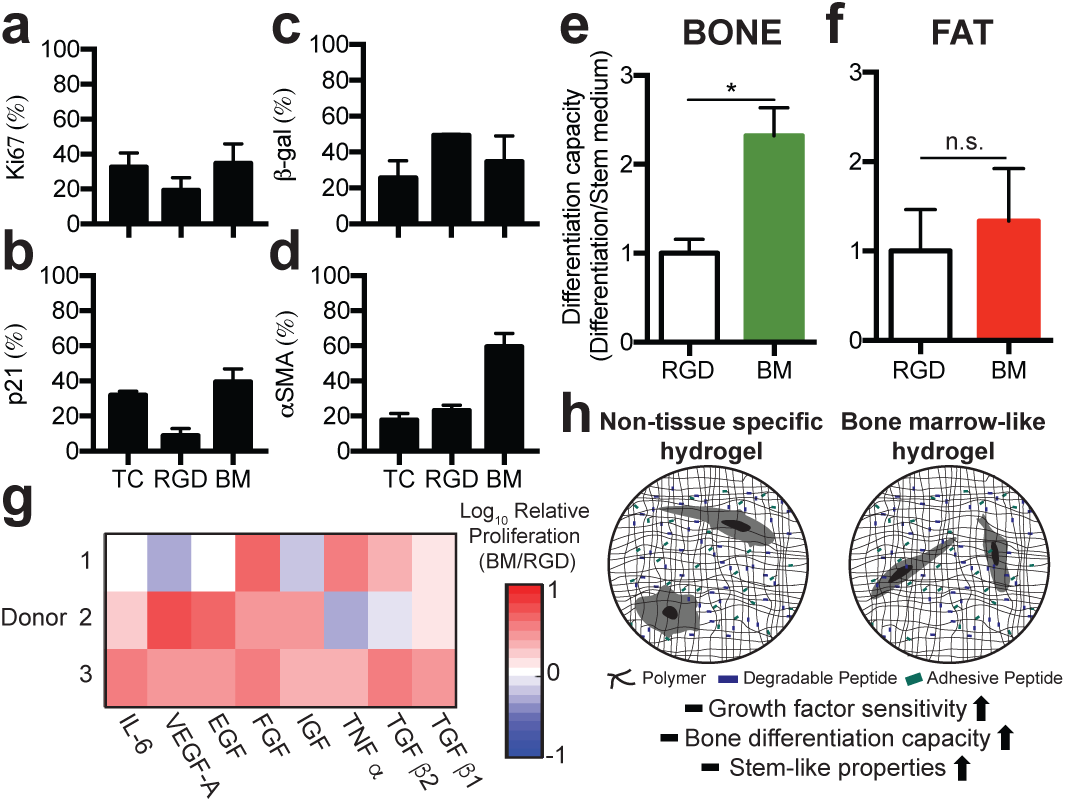
The bone marrow hydrogel maintains MSC stemness. Staining for a) Ki67, b) p21, c) beta-galactosidase, and d) α-smooth muscle actin positive cells in a hydrogel with no degradability and 2 mM RGD (RGD) or the bone marrow hydrogel (BM). e) Oil Red O or f) Osteoimage differentiation capacity normalized to the RGD hydrogel. g) Log_10_ of cell metabolic activity three days after cell encapsulation into the bone marrow hydrogel or an RGD hydrogel for all donor MSCs. Each growth factor was dosed at 20 ng/mL in cell culture medium. h) Schematic to compare how the two hydrogels impact observed MSC phenotypes.

We next explored whether MSCs were differentiating. Interestingly, α-smooth muscle actin was highest in the bone marrow hydrogel, suggesting reduced clonogenicity and fat differentiation (Figure 5d)^38,39^. All donor MSCs were capable of differentiating into bone and fat, shown by staining hydroxyapatite or lipids, respectively (Suppl. Figure 5). Differentiation capacity was measured by quantifying the ability of cells to differentiate in the presence or absence of differentiation medium. In the bone marrow hydrogel, MSCs had a higher capacity to differentiate into bone compared to RGD-functionalized hydrogels (Figure 5e). In both the RGD-functionalized and bone marrow hydrogels, spontaneous hydroxyapatite formation was observed without the presence of differentiation cues (Suppl. Figure 6). Adipose differentiation was similar in both materials (Figure 5f). Our results correlate with reports that α-smooth muscle actin positive MSCs filtered from bone marrow have a higher osteogenic differentiation potential^39^. Interestingly, this study also found that these cells had a higher *in vivo* tissue regeneration capacity.

This led us to hypothesize that the bone marrow hydrogel provided a niche for MSCs to differentiate and respond to growth factors typically present in the bone milieu that are responsible for MSC activation, differentiation, proliferation, and trafficking^40,41^. We treated MSCs encapsulated in hydrogels with a panel of proteins associated with either MSC differentiation or proliferation^42^. We observed that MSCs encapsulated in the bone marrow hydrogel were more metabolically active when exposed to this panel than when encapsulated in the RGD-functionalized hydrogel (Figure 5g). This supports the notion that specific integrin binding in the marrow influences soluble factor signaling.

The described 3D bone marrow hydrogel is highly tunable and can be readily used to probe the underlying mechanisms driving cellular differentiation and phenotypes caused by soluble cues. *In vivo* the bone marrow niche needs to be able to support progenitor populations and to direct cell differentiation, a feature we demonstrate here. Overall, we only observed this unique biological response by combining both the physical and chemical properties of real bone marrow into a PEG hydrogel. While we have focused specifically on bone marrow, this approach could be applied to any tissue in the human body, and represents the next generation of tissue-driven hydrogel design for cell culture.

## Outlook

Here, we combined proteomic-based bioinformatics and biomechanics to make a bone marrow-customized PEG hydrogel. This marrow-mimicking hydrogel is composed simply of PEG and peptides, which polymerizes in 10 seconds under physiological conditions. The novelty of our hydrogel is that it includes 20 unique peptide moieties to more fully capture the integrin-binding and MMP sensitive domains of ECM proteins typical of marrow. Competitor hydrogels typically incorporate 1-3 of these types of peptides and approximate native tissue by incorporating tissue-specific cells. Another approach is to implant bone-like scaffolds into mice to recruit cells and then use *ex vivo* culturing to maintain bone marrow cell populations in culture long-term^43-45^. This latter approach is labor intensive and requires technical expertise to fabricate, limiting its throughput. We also argue that these models under-represent the chemical diversity of native tissue because, while they capture the hierarchical structure of bone, they neglect the unique protein properties present in bone marrow.

Decellularized matrix is currently the only *in vitro* material capable of including the protein complexity of real tissue^46,47^. It is time consuming to make and not batch-controlled, leading to inconsistent results and an inability to separate ensuing variability between the cells and the starting material. Tissue-specific cells can also be made to secrete their own matrix for cell culture use, but this matrix is not necessarily representative of the native environment^48^. As an alternative, we demonstrate an approach to synthetically represent the tissue-specific properties of bone marrow, while maintaining control and simplicity. One appeal of this system is that it could be used to co-culture cells or be formed around any cell or organoid of interest^11^. Additionally, because features can easily be tuned, pseudo ECM knock-out environments can be used to understand ECM-mediated cell signaling. Future work should focus on a more thorough understanding of how each component of the ECM, and how perturbations of these properties, contributes to and changes observed cell phenotypes. In sum, we have captured the ECM of real bone marrow using simple chemistry in a widely-used material that is adaptable to high throughput, systems-level screens^49^. We propose this approach could be applied to any tissue or organ, creating a new class of designer biomaterials that can be employed to elucidate ECM-driven mechanisms in cells not easily achieved by other systems.

## Materials and methods

### Cell culture

All cell culture supplies were purchased from Thermo Fisher Scientific (Waltham, MA) unless otherwise noted. Human mesenchymal stem cells (MSCs) were received through a material transfer agreement with Texas A&M University College of Medicine Institute for Regenerative Medicine at the Scott & White Hospital funded by the National Institute of Health (NIH). MSCs from three donors were cultured in alpha minimum essential medium (αMEM), supplemented with 16.5% fetal bovine serum (FBS) and 1% L-glutamine, and used between the 2^nd^ and 6^th^ passage. The hTERT MSCs were provided from Dr. Junya Toguchida and the human breast cancer cell line MDA-MB-231 was provided by Dr. Shannon Hughes. These were cultured in Dulbecco’s modified eagle’s medium (DMEM), supplemented with 1% L-glutamine, 1% penicillin– streptomycin, 10% FBS, 1% non-essential amino acids, and 1% sodium pyruvate.

### Identifying integrin-binding and MMP-degradable proteins in bone marrow

Manual data mining was used to identify 48 integrin-binding proteins and 44 MMP-degradable proteins (Suppl. Table 1 and 2). These proteins were quantified in human bone marrow using the Protein Atlas (Suppl. Table 3)^8^. The histological score was annotated for each protein. The value of the histological score for the hematopoietic cells was averaged across all the patients scored. This list was used to identify which proteins or protein substrates would be represented by integrin-binding moieties or degradable peptide sequences for the majority of the proteins identified in bone marrow tissue. The histological value was used to determine the percentage of each integrin-binding peptide and MMP-degradable crosslinker to use for proteins in bone marrow.

### Solid-phase peptide synthesis

All peptides were synthesized on a CEM’s Liberty Blue automated solid phase peptide synthesizer (CEM, Mathews, NC) using Fmoc protected amino acids (Iris Biotech GMBH, Germany). Peptide was cleaved from the resin by sparging-nitrogen gas through a solution of peptide-resin and trifluoroacetic acid (TFA), triisopropylsilane, water, and 2,2′- (Ethylenedioxy)diethanethiol at a ratio of 92.5:2.5:2.5:2.5 % by volume, respectively (Sigma-Aldrich, St. Louis, MO) for 3 hours at room temperature in a peptide synthesis vessel (ChemGlass, Vineland, NJ). The peptide solution was filtered to remove the resin and the peptide was precipitated out using diethyl ether at -80°C (Thermo). Molecular mass was validated using a MicroFlex MALDI-TOF (Bruker, Billerica, MA) using α-cyano-4-hydroxycinnamic acid as the matrix (Sigma-Aldrich). Peptides were purified to ≥95% on a VYDAC reversed-phase c18 column attached to a Waters 2487 dual **λ** adsorbable detector and 1525 binary HPLC pump (Waters, Milford, MA).

The following sequences were synthesized: GCGDGEA, GPRGGC, CSRARKQAASIKVAVADR, CSVTCG, CGGYSMKKTTMKIIPFNRLTIG, GCKQLREQ, GCDPGYIGSR, GRGDSPCG, GCRDRPFSMIMGDRCG, GCRDGPLGLWARDRCG, GCRDVPLSLTMGDRCG, GCRDGPQGIWGQDRCG.

The following sequences were purchased from GenScript (China) at >96% purity: CGGSVVYGLR, CGPHSRNGGGGGGRGDS, CGP(GPP)_5_GFOGER(GPP)_5_, CGGAEIDGIEL, GCRDIPESLRAGDRCG, GCGGQWRDTWARRLRKFQQREKKGKCRKA, GCRDVPLSLYSGDRCG, GCRDSGESPAYYTADRCG, GCRDVPMSMRGGDRCG

### Polymerization of 3D bone marrow hydrogels

A 20K 8-arm PEG-maleimide (Jenkem Technology, Plano, TX) was reacted with 2mM of the bone marrow integrin-binding peptide cocktail (Suppl. Table 4) for 10 minutes in serum free medium at pH 7.4. This solution was crosslinked at a 1:1 molar ratio of thiol to maleimide in PBS at pH 7.4, and the crosslinker cocktail was composed of 75 molar% of 1.5K linear PEG-dithiol (Jenkem) and 25 molar% of the MMP-degradable cocktail (Suppl. Table 5). Gels were polymerized in 10 μL volumes with 1,000 cells/μL and cell culture medium was added after 5 minutes to swell the material for at least 18 hours before use. Other hydrogel combinations were made with a 2K, 10K, and 20K 4-arm PEG-maleimide, all crosslinked at a 1:1 molar ratio of thiol to maleimide with 1.5K linear PEG-dithiol.

### ECM protein enrichment from tissues

Tissue samples from healthy women between ages 45-60 were obtained from Cooperative Human Tissue Network funded by the National Cancer Institute (NCI) under IRB exempt status. Insoluble ECM proteins were extracted from 500 mg of tissue using the CNMCS compartmental protein extraction kit according to the manufacturer’s instructions (Millipore, Billerica, MA). This resulted in an insoluble ECM pellet.

### Mass spectrometry

Two biological replicates were analyzed for human bone marrow, brain, and lung tissues. The ECM-rich pellet remaining from the CNCMS kit was solubilized and reduced in 8 M urea, 100 mM of ammonium bicarbonate, and 10 mM dithiothreitol (DTT) (Thermo) for 30 minutes at pH 8 and 37°C. Samples were alkylated with 25 mM iodoacetamide (Sigma-Aldrich) in the dark at room temperature for 30 minutes before the solution was quenched with 5 mM DTT. Prior to cleavage, the solution was diluted to 2 M urea with 100 mM ammonium bicarbonate at pH 8. Proteins were cleaved via trypsin (Thermo) and Lys-C endoproteinase (Promega, Madison, WI), at a ratio of 1:50 enzyme to protein overnight (12-16 hours) at 37°C. Samples were cleaned and concentrated using a C18 column (Thermo). A reverse phase LC gradient was used to separate peptides prior to mass analysis. Mass spectrometry analysis was performed in an Orbitrap Fusion Tribrid (Thermo). Peptides were aligned against the Matrisome using the Thermo Proteome Discoverer 1.41.14^18^. Parameters used trypsin as a protease, with 4 missed cleavage per peptide, a precursor mass tolerance of 10 ppm, and fragment tolerance of 0.6 Da.

### MMP degradation of bone marrow tissue

The MMP degradation assay was adapted from a protocol by Skjøt-Arkil et al. [6]. The ECM-rich pellet from the CNMCS kit was solubilized in 8 M urea at pH 8 and lyophilized in 200 μg aliquots. The lyophilized ECM was resuspended in 100 mM Tris-HCl, 100 mM NaCl, 10 mM CaCl_2_, and 2 mM ZnOAc at pH 8.0. (Sigma-Aldrich) MMP 1, MMP-3 (901-MP, 513-MP, R&D Systems, Minneapolis, MN) MMP 2, MMP 9, MMP 13, MMP 14 (ab125181, ab168863, ab134452, ab168081, Abcam, Cambridge, MA), and MMP 7 (CC1059, Millipore) were activated according to the manufacturer’s instructions and mixed individually with 200 μg of tissue per 1 μg of either active enzyme, or MMP buffer was used as a control. Samples were mixed for 18 hours at 37°C, at which point the reaction was terminated with 25 μM of GM6001 (Millipore). Digested protein was run on a Novex 12% Tris-glycine polyacrylamide gel, stained using silver stain (Thermo) and imaged using the IN Genius Syngene Bioimaging platform (Frederick, MD).

### Competitive binding assay

Glass coverslips were prepared with 1 ug/cm^2^ of the bone marrow peptide coupled to the surface using amine chemistry described by Barney et al^2^. Cells were seeded at 4,000 cells/cm^2^ in their normal growth medium after 30 minutes of pretreatment with individual peptides or the complete bone marrow cocktail. Bone marrow was dosed at a molar amount of 25 nmol/mL of medium and the molar amount dosed for each individual peptide was as follows: GRGDSPCG at 600 pmol/mL, CGPHSRNGGGGGGRGDS and GCGGQWRDTWARRLRKFQQREKKGKCRKA at 220 pmol/mL, CGP(GPP)_5_GFOGER(GPP)_5_, CGGSVVYGLR, and GPRGGC at 160 pmol/mL, CSVTCG and CGGYSMKKTTMKIIPFNRLTIG at 100 pmol/mL, GCGDGEA, CSRARKQAASIKVAVADR, GCKQLREQ, and CGGAEIDGIEL at 60 pmol/mL, and GCDPGYIGSR at 40 pmol/mL. Cells were imaged beginning 10 minutes after seeding in an environment controlled Zeiss Axio Observer Z1 microscope (Carl Zeiss, Oberkochen, Germany) using an AxioCam MRm camera and an EC Plan-Neofluar 20X 0.4 NA air objective. Images were taken using Zeiss Axio Observer Z1 (Carl Zeiss) at five-minute intervals for 2 hours and cell areas were traced in ImageJ (NIH, Bethesda, MD).

### Outgrowth of cells into MMP sensitive hydrogels

Cytodex1 microcarrier beads (Sigma-Aldrich) were swollen in sterile 1X PBS (1 g beads/50 mL PBS) and autoclaved for 30 minutes at 121°C. Flasks were coated with poly (2-hydroxyethy methacrylate) suspended in ethanol at 20 mg/mL and allowed to evaporate in a biosafety cabinet for 30 minutes to make them non-adherent. Cells were seeded at 10-50 cells/bead in non-adherent flasks at a 0.1 mL of beads/mL of media. The flask was shaken every hour for 4 hours to ensure coating onto beads, and cells were allowed to grow on beads for 48 hours post-seeding. Hydrogels were prepared with 4-arm PEG-maleimide at a 20wt% cross-linked at a 1:1 molar ratio with 50% 1.5K linear PEG-dithiol and 50% of each individual MMP degradable peptide sequence (Suppl. Table 5). Hydrogels were imaged at days 1, 3, and 6 and all image analysis was performed in ImageJ (NIH).

### Validation of peptide incorporation

The Measure-iT thiol kit was used to quantify unreacted thiols (Thermo). Buffers were prepared according to the manufacturer’s guidelines. Mono-functional peptides were incorporated at 1 mM in a 100 μL volume of PEG-maleimide for 10 minutes before reacting with 100 μL of the Measure-iT thiol working solution. Di-functional peptides were reacted with PEG-maleimide in 10 μL volumes for 10 minutes before reacting with 100 μL of the Measure-iT thiol working solution. The hydrogel was reduced by immersing hydrogels in sodium borohydride (NaBH, Sigma-Aldrich) in water at a molar ratio of 4:1 NaBH to thiol for 4 hours before adding Measure-iT thiol working solution. All solutions or hydrogel supernatants were read at an excitation of 494 nm and emission of 517 nm within 5 minutes of the reaction. To quantify which peptides did not react, the supernatant from a hydrogel swollen in water for 2 hours was lyophilized, resuspended in 1:1 acetonitrile and ultrapure water with 0.1% TFA at a theoretical concentration 100 pmol/μL, assuming 0% of the peptides coupled to the hydrogel. Peptides were identified using a MicroFlex MALDI-TOF (Buker) with either saturated α-cyano-4-hydroxy cinnamic acid or 10 mg/mL 2,5-dihydroxybenzoic acid as our matrix (Sigma-Aldrich).

### Hydrogel mechanical and structural characterization

The effective Young’s modulus was measured using indentation testing on 10 μL volumes of the 3D hydrogels. A custom-built instrument was used as previously described^50^. Bone marrow mechanical data was taken from Jansen et al^7^. For this application, a flat punch probe was applied to samples at a fixed displacement rate of 10 μm/s, for maximum displacement of 100 μm. The first 10% of the linear region of the force-indentation curves were analyzed using a Hertzian model modified by Hutchens et al. to account for dimensional confinement described by the ratio between the contact radius (a) and the sample height (h) (0.5<a/h<2)^51^.

### MSC spreading with varying peptide concentrations

hTERT MSCs were encapsulated into the 3D bone marrow hydrogels (described above) with peptide concentrations varying from 0 to 4 mM of the bone marrow peptide cocktail. After 24 hours, hydrogels were fixed in 10% formalin for 10 minutes and stained with AlexaFluor 555 phalloidin (A34055, ThermoFisher, 1:40) and DAPI (ThermoFisher, 1:10,000). Cells were imaged Zeiss Spinning Disc Observer Z1 microscope (Carl Zeiss) using an HRm AxioCam and an EC Plan-Neofluar 20X 0.5 NA air objective. Images were taken using Zen (Carl Zeiss) and cell areas were traced in ImageJ (NIH).

### Differentiation of MSCs across biomaterials

Differentiation of cells was assayed across 5 different biomaterial platforms: tissue culture polystyrene, glass coverslips, 2D PEG hydrogels, and 3D PEG hydrogels with either the bone marrow cocktail or the RGD peptide functionality. Glass coverslips were prepared the same way as for the competitive binding assay. 2D PEG-phosphorylcholine (PEG-PC, Sigma-Aldrich) hydrogels were prepared with bone marrow peptides coupled to the surface at 1 ug/cm^2^ as described by Herrick et al^3^. PC was kept at 17 wt% (0.6 M) and PEG is added at 1.1 wt% (0.015 M) for a ∼4 kPa hydrogel. Cells were seeded at a density of 15,000 cells/cm^2^ on plastic and coverslips, 30,000 cells/cm^2^ for 2D hydrogels, and at a density of 2,000 cells/μL in 3D platforms. For osteoblast differentiation, cells were provided cell culture medium supplemented with 10 mM glycerol phosphate (Santa Cruz Biotechnology, Dallas, TX), 1 nM dexamethasone, and 50 M L-ascorbic acid 2-phosphate (Sigma-Aldrich). For adipose cell differentiation, cells were provided cell culture medium supplemented with 0.5 μM isobutylmethylxanthine, 0.5 M dexamethasone, and 50 M indomethacin (Sigma-Aldrich). Cells were maintained for 21 days with medium changes every 3-4 days. After 21 days, cells and materials were fixed in 10% formalin (Thermo) prior to staining. Oil Red O (Thermo) staining was used to identify lipid formation and hydroxyapatite formation was identified using an Osteoimage mineralization assay (Lonza, Basel, Switzerland). Both staining procedures were performed according to the manufacturer’s instructions. Differentiation capacity was determined by dividing the percentage of cells that differentiated in differentiation medium by the percentage that differentiated in stem cell medium. This number for both conditions was normalized to the RGD hydrogel.

### Cell proliferation in response to growth factors

MSCs were seeded at 1,000 cells/μL in the bone marrow hydrogel or a 20wt%, 20K PEG-maleimide functionalized with 1 mM GRGDSPC (Genscript) crosslinked 100% with 1.5K PEG-dithiol. Gels were individually dosed with 20 ng/mL of select growth factors: transforming growth factor- β_1_ (Millipore), transforming growth factor-β_2_ (Sigma-Aldrich), transforming growth factor-α, insulin-like growth factor, fibroblast growth factor-1, epidermal growth factor (R&D Systems), vascular endothelial growth factor-A, and interleukin-6 (Abcam). After 5 days in culture, with media changes every 2 days, cell proliferation was measured with CellTiter 96 AQueous One Solution Cell Proliferation Assay (Promega) at 490 nm (BioTek ELx800 microplate reader, Winooski, VT). Final results were normalized to a proliferation reading of cells grown in hydrogels for 24 hours in normal cell culture medium.

### Immunofluorescence and senescence stains

After 7 days, cells were fixed, permeabilized, and stained. The following antibodies were used for immunofluorescence: Ki67 (ab16667, 1:200, Abcam), p21 (ab7903, 1:200, Abcam), alpha smooth muscle actin (ab7817, 1:200, Abcam). Beta-galactosidase activity was determined using the Senescence Cell Histochemical Staining Kit (Sigma-Aldrich) according to the manufacturer’s instructions. Cell nuclei were stained with DAPI at a 1:10,000 dilution. Samples were imaged on a Zeiss Cell Observer SD (Carl Zeiss).

### Statistical Analysis

Statistical analysis was accomplished using GraphPad’s Prism v7.0a. Data is reported as the mean ± standard error. Unless otherwise noted, a two-tailed t-test was used. P-values <0.05 are considered significant, where p<0.05 is denoted with *, ≤0.01 with **, ≤0.001 with ***, and ≤0.0001 with ****.

## Acknowledgements

We would like to thank Dr. Sarah Perry, Dr. Peter Chien, and Dr. Lila Gierasch for technical assistance and use of equipment. Research reported in this publication was supported by the Office of the Director, National Institutes of Health of the National Institutes of Health under Award Number S10OD010645. SRP is a Pew Biomedical Scholar supported by the Pew Charitable Trusts. SRP was supported by a faculty development award from Barry and Afsaneh Siadat. This work was funded by an NIH New Innovator award (1DP2CA186573-01) and a National Science Foundation (NSF) CAREER grant (DMR-1454806) to SRP.

## Author contributions

LEJ analyzed, designed, and performed all experiments. TPM assisted with peptide synthesis and characterization. SRP supervised the study. LEJ and SRP wrote the manuscript.

## Competing financial interests

The authors declare no competing financial interests.

